# Loss of SHMT2 mediates 5-FU chemoresistance by inducing autophagy in colorectal cancer

**DOI:** 10.1101/680892

**Authors:** Jian Chen, Guangjian Fan, Chao Xiao, Xiao Wang, Yupeng Wang, Guohe Song, Xueni Liu, Jiayi Chen, Huijun Lu, Weiping Guo, Huamei Tang, Guohong Zhuang, Zhihai Peng

**Author notes:** **Corresponding authors**: **Zhihai Peng, *M.D. & Ph. D.*** Cancer Research Center of Xiamen University, Department of General Surgery, Xiang’an Hospital of Xiamen University, School of Medicine, Xiamen University, 2000 Xiang An East Road, Xiamen 361100, China.; **Guohong Zhuang, *M.D. & Ph. D.*** Organ Transplantation Institute of Xiamen University, Fujian Provincial Key Laboratory of Organ and Tissue Regeneration, School of Medicine, Xiamen University, 4221-122 Xiang An South Road, Xiamen 361102, China.; **Huamei Tang, *M.D. & Ph. D.*** Cancer Research Center of Xiamen University, Department of General Surgery, Xiang’an Hospital of Xiamen University, School of Medicine, Xiamen University, 2000 Xiang An East Road, Xiamen 361100, China.; **Weiping Guo, *M.D. & Ph. D.*** Department of Gastrointestinal Surgery, The Third Affiliated Hospital of Sun Yat-Sen University, 600 Tianhe Road, Guangzhou 510630, China. These authors contributed equally to this work (Jian Chen, Guangjian Fan, Chao Xiao, Xiao Wang).

## Abstract

Serine hydroxymethyltransferase 2 (SHMT2) plays a vital role in one-carbon metabolism and drives colorectal carcinogenesis. In our study, loss of SHMT2 induced 5-Fluorouracil (5-FU) chemoresistance and was associated with poor prognosis in colorectal cancer (CRC). To elucidate the possible mechanism and generate a strategy to sensitize CRC to 5-FU chemotherapy, we first identified the binding proteins of SHMT2 in cancer cells by mass spectrometry. We found that SHMT2 inhibited autophagy through binding cytoplasmic p53. In fact, SHMT2 prevented cytoplasmic p53 degradation by inhibiting the binding of p53 and HDM2. Under treatment with 5-FU, depletion of SHMT2 promoted autophagy and inhibited apoptosis. Autophagy inhibitors CQ decreased low SHMT2-induced 5-FU resistance in vitro and in vivo. Finally, we enhanced the lethality of 5-FU treatment to CRC cells through the autophagy inhibitor or knockdown of SHMT2 in patient-derived and CRC cell xenograft models. Our findings identified the low SHMT2-induced autophagy on 5-FU resistance in CRC. These results reveal SHMT2-p53 as a novel cancer therapeutic target to reduce chemotherapeutic drug resistance.

## Introduction

Colorectal cancer (CRC) is the third leading cause of cancer mortality worldwide due to its metastatic properties and resistance to current treatment (1). 5-Fluorouracil (5-FU)-based systemic adjuvant chemotherapy is the widely accepted systemic therapy option for CRC patients, highlighting the need to better understand the underlying chemoresistance mechanism and identify tumour cell-specific therapeutic targets for drug discovery or “repositioning” of known therapies (2,3).

Public microarray and RNA-seq data from the GEO and TCGA databases provide information about the expression of multiple genes in CRC. By deep learning methods, we found that MET, CPM, SHMT2, GUCA2B, and SCN9A are significantly related to CRC progression. Among these genes, serine hydroxymethyltransferase 2 (SHMT2) plays a regulatory role in the conversion of serine to glycine, which controls cell proliferation (4,5). SHMT2 is a potential cancer driver gene and to promote colorectal carcinogenesis (6,7). SIRT3 and SIRT5 deacetylate SHMT2 at Lys 95 and 280, respectively, to increase enzymatic activity and drive cancer cell proliferation (7,8). Additionally, SHMT2 functions as a component of BRISC-SHMT2 complexes to deubiquitinate the type 1 interferon (IFN) receptor chain 1 (IFNAR1) and HIV-1 Tat in the cytoplasm (9,10). Thus, SHMT2 not only is a methyltransferase but also plays a role as a binding protein influencing the degradation of other proteins.

CRC cells with rapid proliferation need to transform moderate amounts of glycine to serine to support nucleotide biosynthesis and tumour cell proliferation (11–16). Thus, SHMT2 is upregulated in cancers to support tumour cell proliferation (6,17–19). SHMT2 is required for glioma cell survival but also renders these cells sensitive to glycine cleavage system inhibition (15). This finding implies that the role of SHMT2 in tumour therapy is complicated and needs to be investigated further.

Autophagy, a catabolic process, is thought to buffer metabolic stress, thereby aiding cell survival (20,21). In the context of cancer, autophagy plays a puzzling role, serving as a tumour suppressor during the initial stages but later protecting tumour cells from chemotherapy and radiotherapy resistance, hypoxia and the immune defense system (22–27). Multiple cellular stressors, including activation of the tumour suppressor p53, can stimulate autophagy (20). Pharmacological inhibition, knockout or knockdown of p53 can induce autophagy (28,29). Cytoplasmic but not nuclear p53 is responsible for autophagy inhibition (28,29). However, the upstream signals controlling cytoplasmic p53 are still unknown. How cytoplasmic p53 influences chemotherapy through autophagy also needs to be explored.

In this study, we first discovered a significant correlation between SHMT2 and CRC progression through bioinformatics analysis. Thus, we further explored the function of SHMT2. Strikingly, we found that samples of colorectal cancer (CRC) with low levels of SHMT2 from different patients showed greater 5-FU resistance than samples with higher levels of SHMT2. To explore the underlying mechanism of 5-FU resistance, we investigated whether low levels of SHMT2 significantly induced autophagy while overexpression of SHMT2 inhibited autophagy through binding and degrading cytoplasmic p53. Finally, we found that the autophagy inhibitor chloroquine (CQ) increased the 5-FU chemosensitivity of CRC samples with low levels of SHMT2. Our study reveals the previously unknown function of SHMT2 in autophagy through maintaining the stability of cytoplasmic p53 and suggests a potential strategy for anticancer chemotherapy.

## Results

### Analysis of CRC via high-throughput databases reveals that SHMT2 is pivotal in CRC

High-throughput gene chips are widely used in cancer research. The gene expression database Gene Expression Omnibus (GEO) is one of the world’s largest gene chip databases. We screened 66 differentially expressed genes associated with CRC progression using 5 compound covariate classifiers (diagonal linear discriminant analysis, Bayesian CCP, nearest neighbor, nearest centroid and support vector machines) in 224 colon cancer tissues and 165 adjacent normal tissues from three GEO data sets (GSE9348, GSE44076, and GSE44861) (**Appendix Fig S1**). Cox regression analysis and sensitivity analysis were used to further screen 46 molecular markers related to survival in 532 data sets with prognostic information (GSE14333, GSE17536 and GSE29621 data sets). The random survival forest algorithm was used to screen the five most essential molecules and construct a multifactor regression risk function, as follows: Risk score = −0.370*CPM −0.122*GUCA2B+0.332* MET+0.088* SCN 9A+0.827* SHMT2. These results indicate that these five genes are significantly correlated with CRC progression. Among these genes, SHMT2 was the most prominent in cancer metabolism but less investigated in CRC than the other genes, thus arousing our interest.

### SHMT2 interacts with cytoplasmic p53

SHMT2 is known to play a vital role in one-carbon metabolism and functions as a binding protein influencing the degradation of other proteins (9,10). Thus, we not only investigated the metabolic regulation of SHMT2 but also analyzed its binding proteins using LC-MS/MS analysis (Fig 1A). The results showed that p53 was one of the SHMT2 binding proteins, while KIAA0157, a known binding protein, was also identified (Fig 1B). HCT116 cells that contained wild-type p53 but not mutant p53 were used to verify the binding of p53 and SHMT2. Coimmunoprecipitation experiments with transiently transfected SHMT2 showed that SHMT2 coprecipitated with p53 (Fig 1C). Endogenous p53-SHMT2 binding was also detectable in HCT116 cells (Fig 1D).

**Figure 1.**
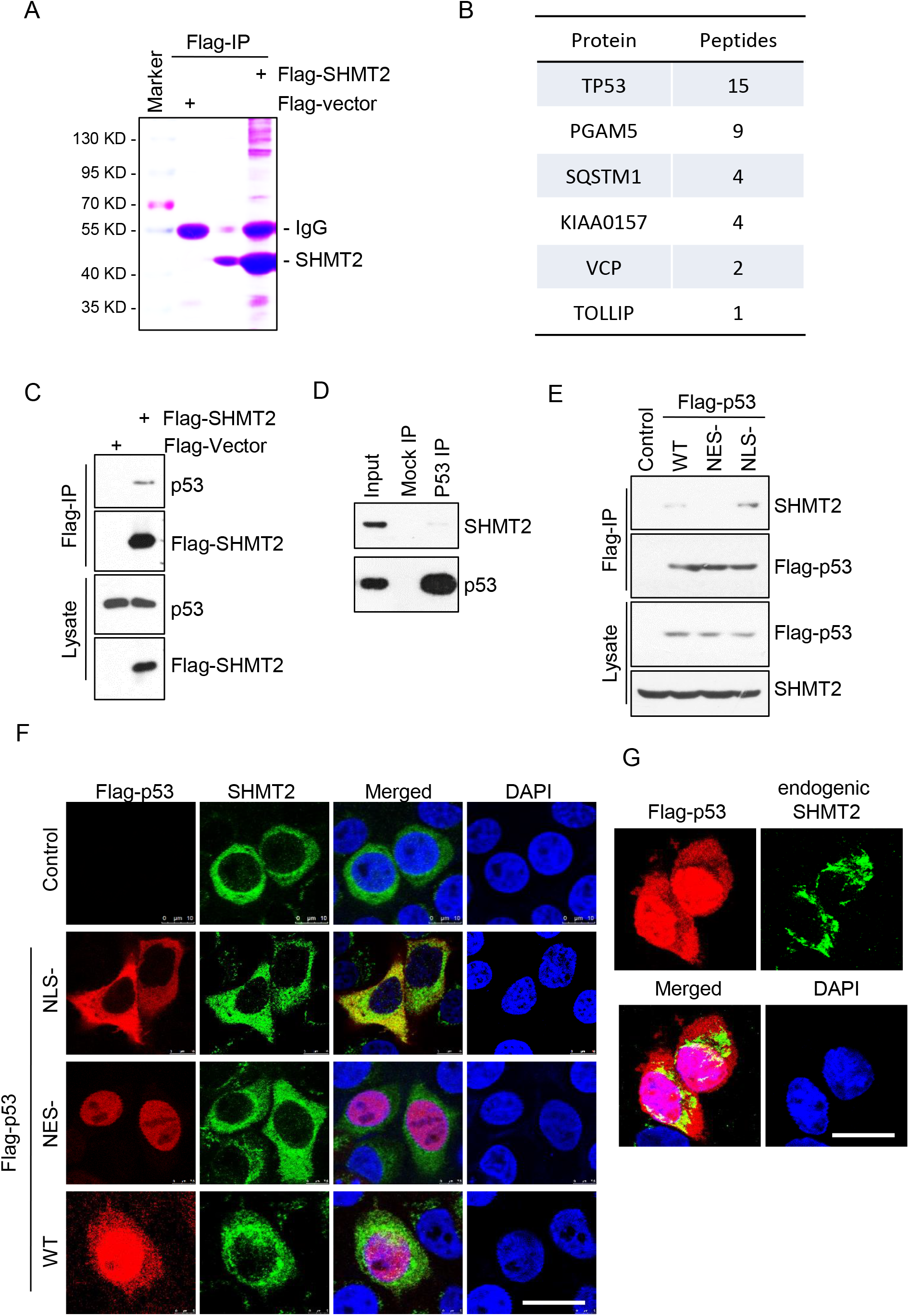
SHMT2 interacts with cytoplasmic p53. A, B SHMT2 purified by Flag-IP was collected after in-gel digestion and used in LC-MS/MS analysis to search for the binding proteins of SHMT2. (A) Flag-SHMT2 was transfected into 293T cells for 24 h, isolated by coimmunoprecipitation, separated by SDS-PAGE and stained using Coomassie. (B) Tabular display of the number of tryptic peptides from each of the indicated proteins that copurified with SHMT2. C HCT116 cells transfected with Flag-SHMT2 were immunoprecipitated with FLAG-M2 beads, followed by western blotting for p53 and SHMT2. D Immunoprecipitation using an anti-p53 antibody (Do-1) was followed by western blotting with anti-SHMT2 or anti-p53 antibodies (ab32389, abcam). E Cytoplasmic p53 bound to SHMT2. HCT116 cells transfected with Flag-WT, nuclear (NES-) or cytoplasmic p53 (NLS-) were immunoprecipitated with FLAG-M2 beads followed by western blotting for FLAG and SHMT2. F, G The colocalization of SHMT2 and cytoplasmic p53. (F) Representative micrographs of HCT116 cells transfected with plasmids expressing WT, nuclear (NES-) and cytoplasmic p53 (NLS-). (G) Representative micrographs of HCT116 cells stained for SHMT2 and p53. Scale bar, 10 μm.

The principal tumour suppressor, p53, accumulates in cells in response to DNA damage, oncogene activation, and other stresses (30). This protein acts as a nuclear transcription factor that transactivates genes involved in apoptosis and numerous other processes (30). However, cytoplasmic p53 was found to inhibit autophagy and trigger apoptosis (28,29,31). Given that SHMT2 locates in mitochondria and the cytoplasm (9), we constructed wild-type p53, cytoplasmic p53 (NLS-), and nuclear p53 (NES-) mutants (29) to identify the correct binding region. Coimmunoprecipitation experiments showed that SHMT2 interacts with cytoplasmic p53 (NLS-) but not nuclear p53 (NES-) (Fig 1E). Moreover, the immunofluorescence results showed that SHMT2 colocalized with cytoplasmic p53 in cells cotransfected with SHMT2, WT p53, nuclear p53 (NES-) or cytoplasmic p53 (NLS-) (29) (Fig 1F). Given that cytoplasmic p53 was also found in mitochondria (31), colocalization of endogenous SHMT2 with cytoplasmic p53 was also detectable (Fig 1G). These results suggest that SHMT2 interacts with cytoplasmic p53.

### Depletion of SHMT2 induces autophagy

Cytoplasmic p53 mediates tonic inhibition of autophagy. Deletion, depletion or pharmacological inhibition of p53 induces autophagy in mouse, human and nematode cells (29). We verified the binding of SHMT2 to cytoplasmic p53, and we then examined the levels of autophagy markers in SHMT2 knockdown (SHMT2-sh) cells. The results showed markedly increased LC3-II levels and decreased p62 levels in SHMT2-sh but not control (scramble-sh) HCT116 cells (Fig 2A). In parallel, SHMT2 knockdown increased the number of LC3 puncta per cell (Fig 2B-C). Transmission electron microscopy analysis of autophagy in SHMT2-sh cells revealed that SHMT2 knockdown led to a marked increase in the number of autophagic vacuoles (AVs) in vitro (Fig 2D-E). Consistent with these findings, increased LC3-II and decreased p62 levels were also observed in SW480 cells (Fig 2F). The number of autophagic vacuoles appreciably increased in SW480 cells (Fig 2G). The above data indicate that SHMT2 inhibits autophagy.

**Figure 2.**
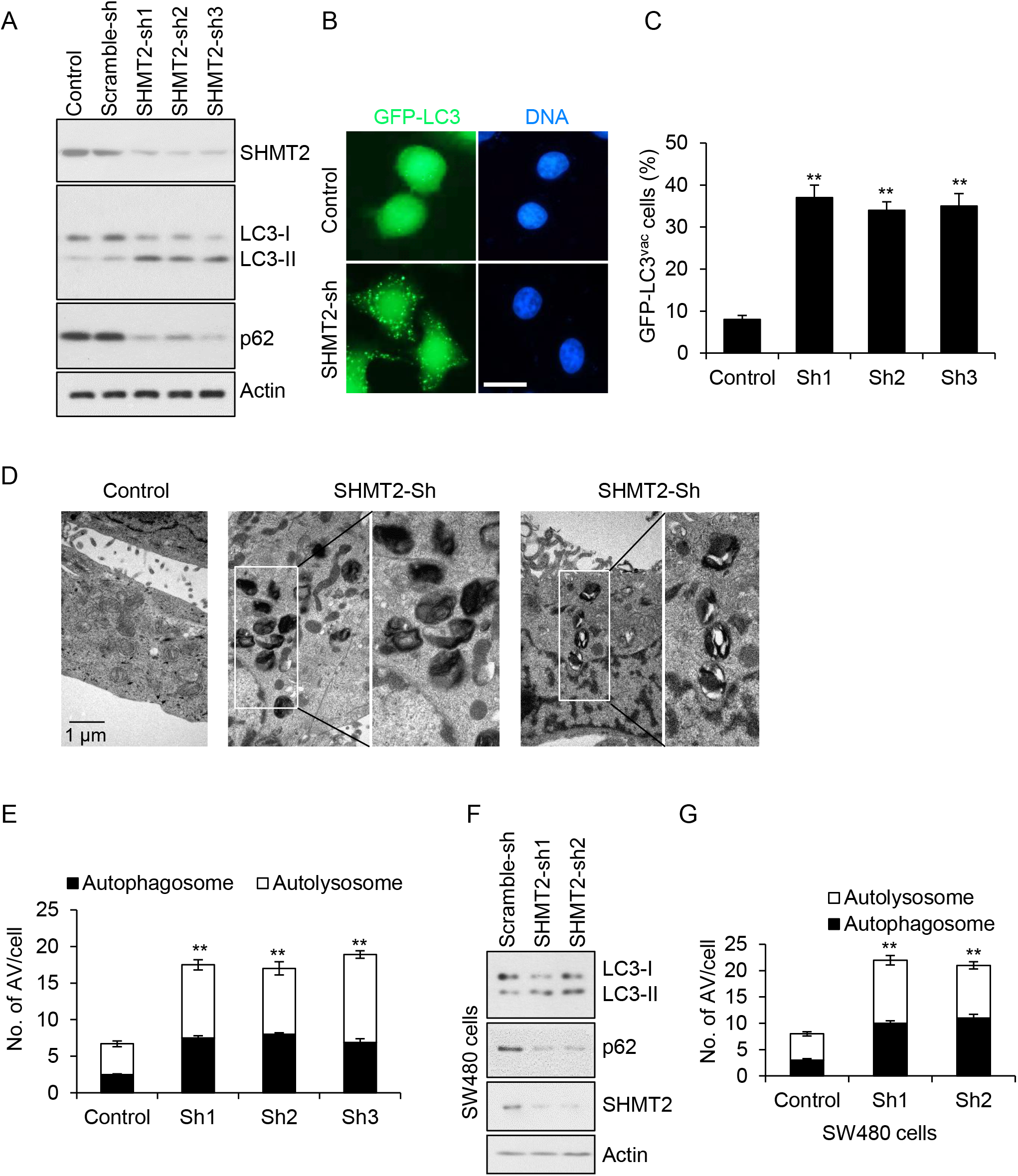
Depletion of SHMT2 induces autophagy. A Effect of SHMT2 on the maturation of LC3. HCT116 cells were infected with Scramble-sh (Control) or SHMT2 knockdown (Sh-1, Sh-2 or Sh-3) lentivirus for 72 h and selected with puromycin (1 mg/ml) to establish stable cell lines. The protein levels of endogenous SHMT2, p62, LC3 and β-actin (as the internal standard) were examined by western blotting using anti-Flag, anti-SHMT2, anti-p62, anti-LC3 and anti-β-actin antibodies, respectively. B GFP–LC3 puncta induced by SHMT2 knockdown. Control and SHMT2-sh stable HCT116 cell lines were transfected with a GFP–LC3 plasmid and cultured in complete medium for 24 hours. Scale bar, 10 μm. C The percentage of cells showing accumulation of GFP–LC3 in puncta (GFP–LC3^vac^) is shown (mean ± s.d., n = 3; \*\**P* < 0.01). D Ultrastructural evidence of autophagic vacuolization induced by SHMT2 depletion. E The number of autophagosomes and autolysosomes was determined for at least 50 cells in 3 independent experiments (mean ± s.d.; \*\**P* < 0.01). F Effect of SHMT2 on the maturation of LC3 in sw480 cells. HCT116 cells were infected with Scramble-sh (Control) or SHMT2 knockdown (Sh-1 or Sh-2) lentivirus for 72 h and selected with puromycin (1 mg/ml) to establish stable cell lines. The protein levels of endogenous SHMT2, p62, LC3 and β-actin (as the internal standard) were examined by western blotting using anti-Flag, anti-SHMT2, anti-p62, anti-LC3 and anti-β-actin antibodies, respectively. G The number of autophagosomes and autolysosomes was determined for at least 50 cells in 3 independent experiments.

### Depletion of SHMT2 induces autophagy through degrading cytoplasmic p53 after 5-FU treatment

Autophagy, as a catabolic process, is thought to buffer metabolic stress, while SHMT2 plays a role as an essential metabolic enzyme to regulate one-carbon metabolism. Thus, we first verified whether SHMT2 regulates autophagy depending on its metabolic regulation. GC-MS (gas chromatography-mass spectrometry) was used to detect metabolites in stably transfected HCT116 cell lines. KEGG enrichment analysis of signaling pathways revealed that SHMT2 might regulate amino acid metabolism (**Appendix Fig S2**). However, no enrichment was found in the regulation of autophagy. This result indicates that SHMT2 may participate in autophagy through other pathways.

As SHMT2 interacts with cytoplasmic p53 (NLS-), we next evaluated whether SHMT2 regulates autophagy via cytoplasmic p53. We transfected a SHMT2 overexpression plasmid into HCT116 cells with p53 deletion (p53-/-) and assessed autophagy. We found that SHMT2 did not affect the LC3II/I ratio or p62 level in the absence of p53 (Fig 3A). Similar results were also observed in p53 knockdown cells (Fig 3B). To further assess whether autophagy inhibition by SHMT2 is p53-dependent, we restored cytoplasmic p53 expression in SHMT2 knockdown cells and found that cytoplasmic p53 reversed the induction of autophagy due to SHMT2 depletion (Fig 3C). Similarly, cytoplasmic p53 also decreased the number of LC3 puncta per cell and the level of GFP-LC3 in SHMT2 knockdown cells (Fig 3D-E). Together, these results indicate that SHMT2 inhibits autophagy via cytoplasmic p53.

**Figure 3.**
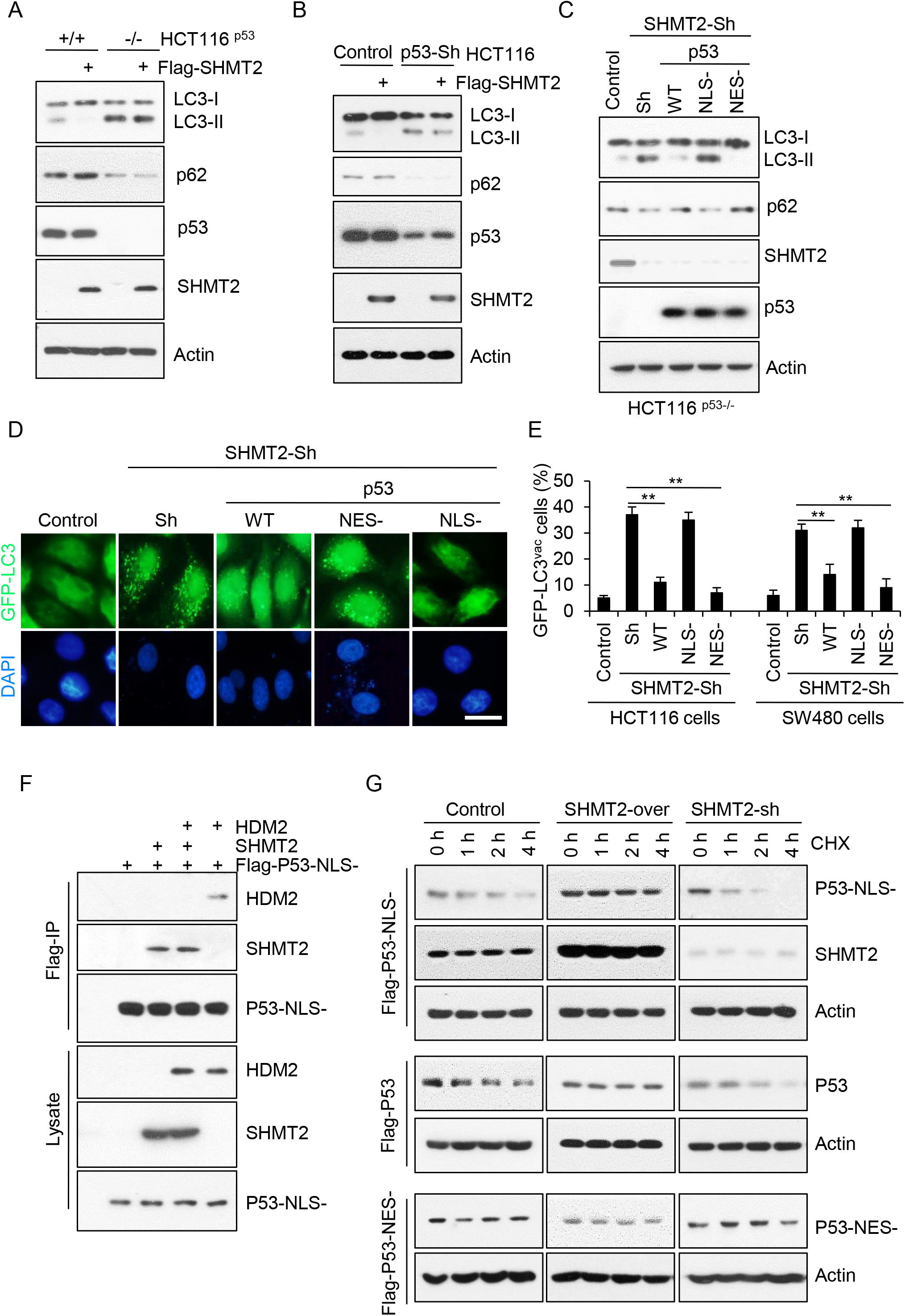
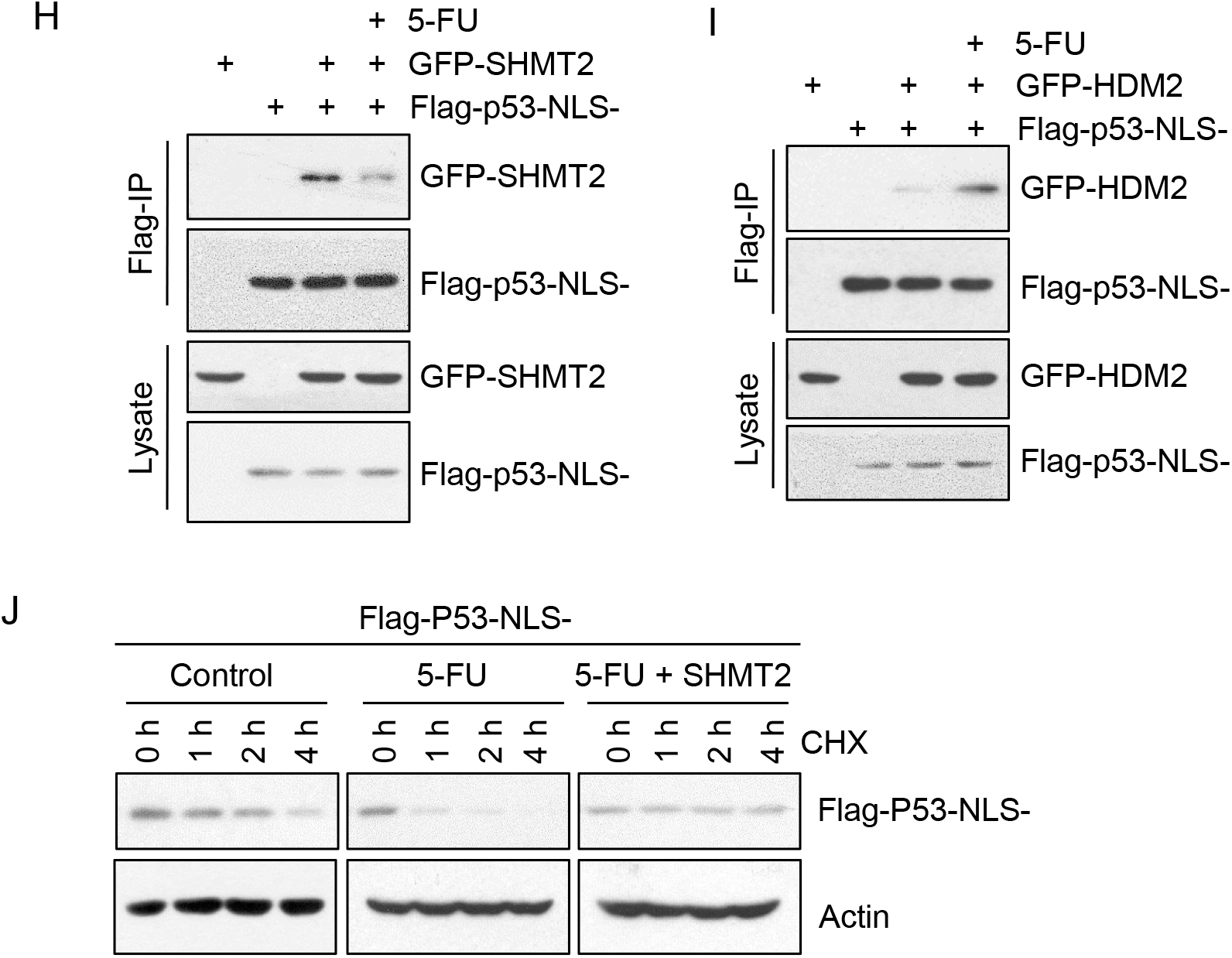
SHMT2 depletion induces autophagy through degrading cytoplasmic p53 in response to 5-FU treatment. A-C Effect of SHMT2 and p53 on the maturation of LC3. The protein levels of SHMT2, p53, p62, LC3 and β-actin (as the internal standard) were assessed by western blotting using anti-Flag, anti-p53, anti-p62, anti-LC3 and anti-β-actin antibodies, respectively. (A) HCT116 cells were infected with Scramble-sh (Control) or p53 knockdown (Sh) lentivirus for 72 h and transfected with Flag-SHMT2 for 24 h. (B) Wild-type (WT) or p53–/– HCT116 cells were transfected with Flag-SHMT2 for 24 h. (C) Control or SHMT2-Sh stable cell lines were transfected with Flag-WT, nuclear (NES-) and cytoplasmic p53 (NLS-) plasmids for 24 h. Scale bar, 10 μm. D GFP–LC3 puncta formation induced by SHMT2-sh or p53 mutants. Control and SHMT2-sh stable HCT116 or SW480 cell lines were transfected with Flag-WT, nuclear (NES-) cytoplasmic p53 (NLS-) or GFP–LC3 plasmids and cultured in complete medium for 24 h. E The percentage of cells showing accumulation of GFP–LC3 in puncta (GFP–LC3^vac^) is shown (mean ± s.d., n = 3; \*\**P* < 0.01). F SHMT2 disrupted the binding of cytoplasmic p53 to HDM2. HCT116 cells transfected with HDM2, SHMT2 and Flag-cytoplasmic p53 (NLS-) were immunoprecipitated with FLAG-M2 beads, followed by western blotting for p53, HDM2 and SHMT2. G SHMT2 maintained the stability of cytoplasmic p53. Western blot analysis of lysates of cell lines with stable SHMT2 overexpression and knockdown transfected with Flag-WT, nuclear (NES-) and cytoplasmic p53 (NLS-) plasmids and treated with the translation inhibitor cycloheximide (CHX, 50 μg/ml) for the indicated hours. H, I HCT116 cells transfected with Flag-cytoplasmic p53 (NLS-), GFP-SHMT2 or GFP-HDM2 were immunoprecipitated with FLAG-M2 beads, followed by western blotting for p53 and GFP. (H) 5-FU disrupted the binding of cytoplasmic p53 to SHMT2. (I) 5-FU promoted the binding of cytoplasmic p53 to HDM2. J Western blot analysis of lysates of HCT116 cell lines transfected with Flag-cytoplasmic p53 (NLS-) or SHMT2 plasmids and treated with CHX for the indicated hours with or without 5-FU.

It was reported that the induction of autophagy stimulates proteasome-mediated degradation of p53 through a pathway dependent on the E3 ubiquitin ligase HDM2 (29,30). Therefore, we investigated whether SHMT2 interferes with p53-HDM2 binding. A coimmunoprecipitation competition assay showed that SHMT2 prevents cytoplasmic p53 from interacting with HDM2 (Fig 3F). Next, we detected the p53 protein level in cells with SHMT2 overexpression or knockdown. The results showed that SHMT2 stabilizes cytoplasmic p53 but not nuclear p53 (Fig 3G). After 5-FU treatment, the binding of SHMT2-p53 decreased (Fig 3H). Conversely, p53-HDM2 binding was increased dramatically upon 5-FU treatment (Fig 3I). As expected, cytoplasmic p53 exhibited rapidly and large-scale degradation under 5-FU treatment, and SHMT2 overexpression significantly decreased the rate of reduction in the p53 protein level (Fig 3J). Taken together, these results indicate that SHMT2 stabilizes p53 by preventing HDM2-mediated degradation in response to 5-FU.

### Inhibition of autophagy induced by low SHMT2 expression sensitizes CRC cells to 5-FU treatment

Given that cytoplasmic p53 triggers apoptosis and inhibits autophagy (31), we further analyzed the balance between apoptosis and autophagy in cells with SHMT2 overexpression and knockdown in response to 5-FU. The levels of cleaved caspase-3, poly (ADP-ribose) polymerase (PARP) and LC3-II were increased in SHMT2-overexpressing cells, while these levels were decreased in SHMT2 knockdown cells, indicating that SHMT2 promotes apoptosis and inhibits autophagy in response to 5-FU (Fig 4A). Autophagy induction prevents tumour cells from undergoing apoptosis and subsequently leads to resistance to chemotherapy (26). To sensitize cells to 5-FU-based chemotherapy, 3-methyladenine (3MA) and chloroquine (CQ), inhibitors of autophagy, were used (25,32). Compared with the control and SHMT2-overexpressing cell lines, SHMT2 knockdown cells showed resistance to 5-FU, and 3-MA or CQ counteracted this resistance (Fig 4B-C).

**Figure 4.**
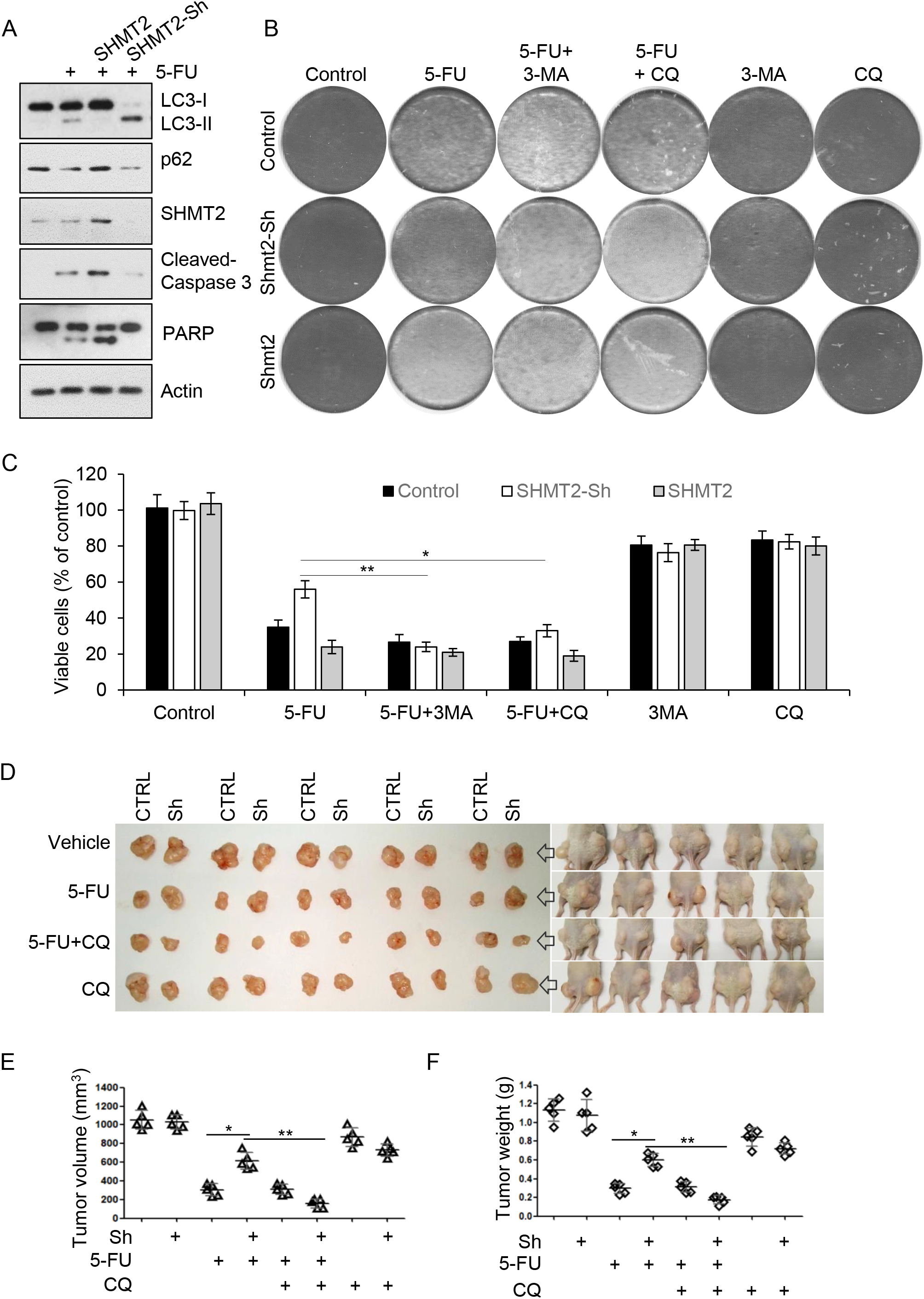
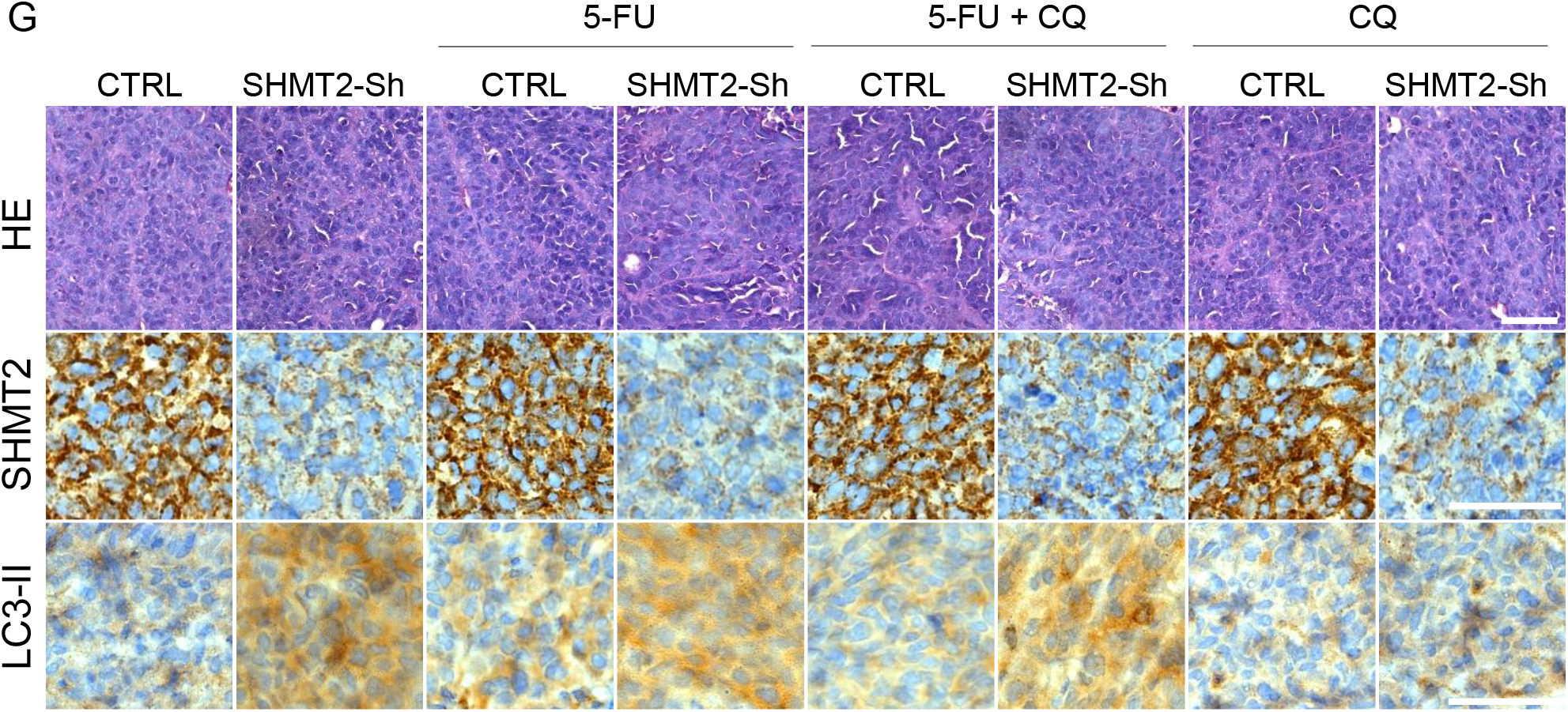
Inhibition of autophagy induced by low SHMT2 expression sensitizes CRC cells to 5-FU treatment. A SHMT2 promotes apoptosis and inhibits autophagy in response to 5-FU. Western blot analysis of lysates of HCT116 cell lines transfected with SHMT2 or infected with SHMT2-Sh virus and treated with 5-FU (10 μM) for 24 h. The protein levels of SHMT2, p62, LC3, cleaved-Caspase 3, PARP and β-actin (as internal standards) were assessed with anti-SHMT2, anti-p62, anti-LC3, anti-cleaved Caspase 3, anti-PARP and anti-β-actin antibodies, respectively. B, C The indicated cells were treated with 5-FU (2 μM), 3-methyladenine (3-MA, 10 mM) or chloroquine diphosphate salt (CQ, 20 μM) for four days and stained with 0.1% crystal violet (B) or analyzed for cell viability by an MTT assay (C), \*\**P* < 0.01. (c-e) The xenograft experiment with Control or SHMT2 knockdown cells treated with 5-FU or CQ is described in the Methods section. D Xenograft tumours were harvested and photographed. E,F The quantification of the (e) average volume and (f) weight of the xenograft tumours is shown. Five tumours from individual mice were included in each group; ** *P* < 0.01. G Sample images of immunohistochemical and hematoxylin-eosin staining (HE) of xenograft tumours. Scale bar, 50 μm.

Next, we investigated whether CQ could enhance the therapeutic response of xenograft tumours with low SHMT2 levels in mice treated with 5-FU. SHMT2 wild-type and SHMT2 knockdown HCT116 cells were injected subcutaneously into nude mice above the left and right hind legs, respectively. The results showed that tumour growth was markedly inhibited in the SHMT2 knockdown groups receiving CQ and 5-FU treatment (Fig 4D-F). These results indicate that the combination of CQ and 5-FU markedly inhibited tumour growth in the SHMT2 knockdown groups. Analysis of tumour tissues by immunohistochemistry (IHC) verified that the expression of LC3-II in the SHMT2 knockdown group with 5-FU treatment increased, while CQ reduced LC3-II expression, indicative of autophagy suppression (Fig 4G). Collectively, our findings show that autophagy induced by low SHMT2 expression led to 5-FU resistance and that inhibition of autophagy sensitized SHMT2-low CRC cells to 5-FU treatment.

### 5-FU resistance is related to low SHMT2 expression and autophagy in human colorectal cancer

SHMT2 is a potential cancer driver gene and to promote colorectal carcinogenesis (6,7), while it is related to 5-FU resistance in CRC cells and xenograft tumours. Thus, we further studied the role of SHMT2 in CRC therapy. q-PCR analysis of 50 paired CRC tissues and adjacent normal tissues showed that SHMT2 expression significantly upregulated in CRC tissues relative to that in normal tissues (**Appendix Fig S3A**). We retrieved the SHMT2 mRNA expression data from the GEO and TCGA databases and found that the expression level of SHMT2 was significantly higher in CRC tissues than in normal mucosa (Figure 5A, **Appendix Fig S3B**).

**Figure 5.**
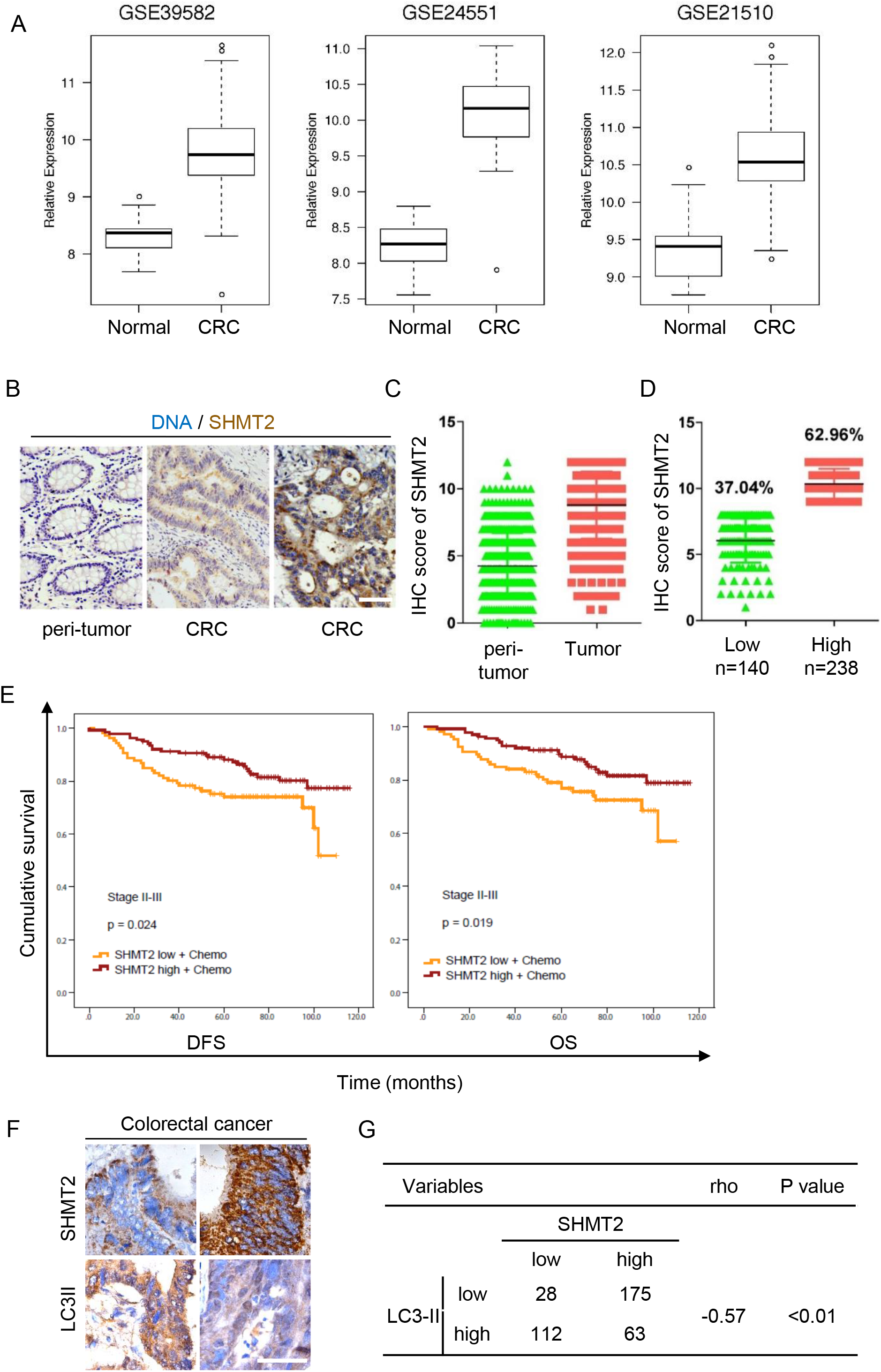
5-FU resistance is related to low SHMT2 expression and autophagy in colorectal cancer. A The expression of SHMT2 in three GEO data sets (GSE39582, GSE24551 and GSE21510). B Representative IHC images of anti-SHMT2 staining in peritumoural and CRC tissues. Scale bar, 50 μm. C A total of 378 paired CRC tissues assessed by IHC are shown. D Among the 378 stage II-III CRC tissues, 37.04% exhibited low expression of SHMT2 (score <9), while 62.96% expressed high expression of SHMT2 (score ≥9). E Survival of patients with different levels of SHMT2 expression. DFS and OS stratified by SHMT2 expression in patients with stage II-III disease and treated with 5-FU–based chemotherapy. F, G Comparison of SHMT2 and LC3-II expression in CRC tissues. (F) Sample images of immunohistochemical staining. Scale bar, 50 μm. (G) Spearman’s r-coefficient test for the evaluation of correlations between the SHMT2 and lc3-II immunohistochemical expression status in CRC tissues. Negative ρ values indicate a negative relationship. The significance level is indicated by the *P* value.

Next, we selected CRC patients with TNM stage II or III disease (n=378) to explore the function of SHMT2 in 5-FU–based adjuvant chemotherapy (**Appendix Table 1**). High expression of SHMT2 was found in human CRC specimens compared to that in normal specimens using immunohistochemical staining (Fig 5B-C). However, considering the complexity of colorectal tumourigenesis, we also found that 37% of CRC tissues showed low SHMT2 expression (Fig 5D). To better understand the contribution of SHMT2 to the prognosis of patients with CRC, especially its effect in response to 5-FU-based adjuvant chemotherapy, we investigated the correlation of DFS (disease-free survival) and OS (overall survival) with SHMT2 expression levels in CRC. Surprisingly, when 5-FU-based adjuvant chemotherapy was administered to patients with SHMT2-low CRC (37%, n=140), these patients showed worse prognoses in terms of DFS and OS (Fig 5E). Moreover, the correlation analysis revealed a significant negative correlation between the protein expression levels of SHMT2 and LC3-II (Spearman’s *ρ*=0.6), indicating that autophagy was induced in SHMT2-low CRC (Fig 5 F-G). Given that the percentage of patients with SHMT2-low CRC is 37%, we need to explore the mechanism underlying 5-FU resistance. These results also imply that the expression of SHMT2 could be used as a marker for 5-FU resistance in clinical therapy.

Taken together, these results further demonstrate that upregulated SHMT2 promotes CRC progression, but low expression of SHMT2 might play an important role in chemoresistance in CRC patients. Furthermore, multivariate Cox proportional hazards analysis suggested that SHMT2 is a new and independent prognostic marker for CRC patients who received 5-FU-based chemotherapy (**Appendix Tables 2 and 3**).

### CQ sensitized xenografts with low SHMT2 expression to 5-FU treatment in the PDX model

The above results showed that low SHMT2 induced 5-FU resistance through autophagy activation. To explore the function of autophagy inhibition in 5-FU therapy, we established a xenograft mouse model in which four CRC patient-derived tissues (two with high expression of SHMT2 and two with low expression of SHMT2) were implanted subcutaneously (Fig 6 A-B). Similar results for LC3-II and p62 expression were observed in xenografts with low expression of SHMT2 as were observed in the abovementioned experiments (Fig 6A). Tumour growth was markedly inhibited in the animals with SHMT2-low xenografts that received combined treatment with CQ and 5-FU compared to that in those receiving 5-FU treatment alone (Fig 6C-E). These results indicate that the combination of CQ and 5-FU markedly inhibited tumour growth in the SHMT2-low group. Analysis of tumour tissues by immunohistochemistry (IHC) verified the expression of SHMT2 and LC3-II in these tumours (Fig 6F). Collectively, these findings show that CQ, as an autophagy inhibitor, sensitized xenografts with low SHMT2 expression to 5-FU treatment.

**Figure 6.**
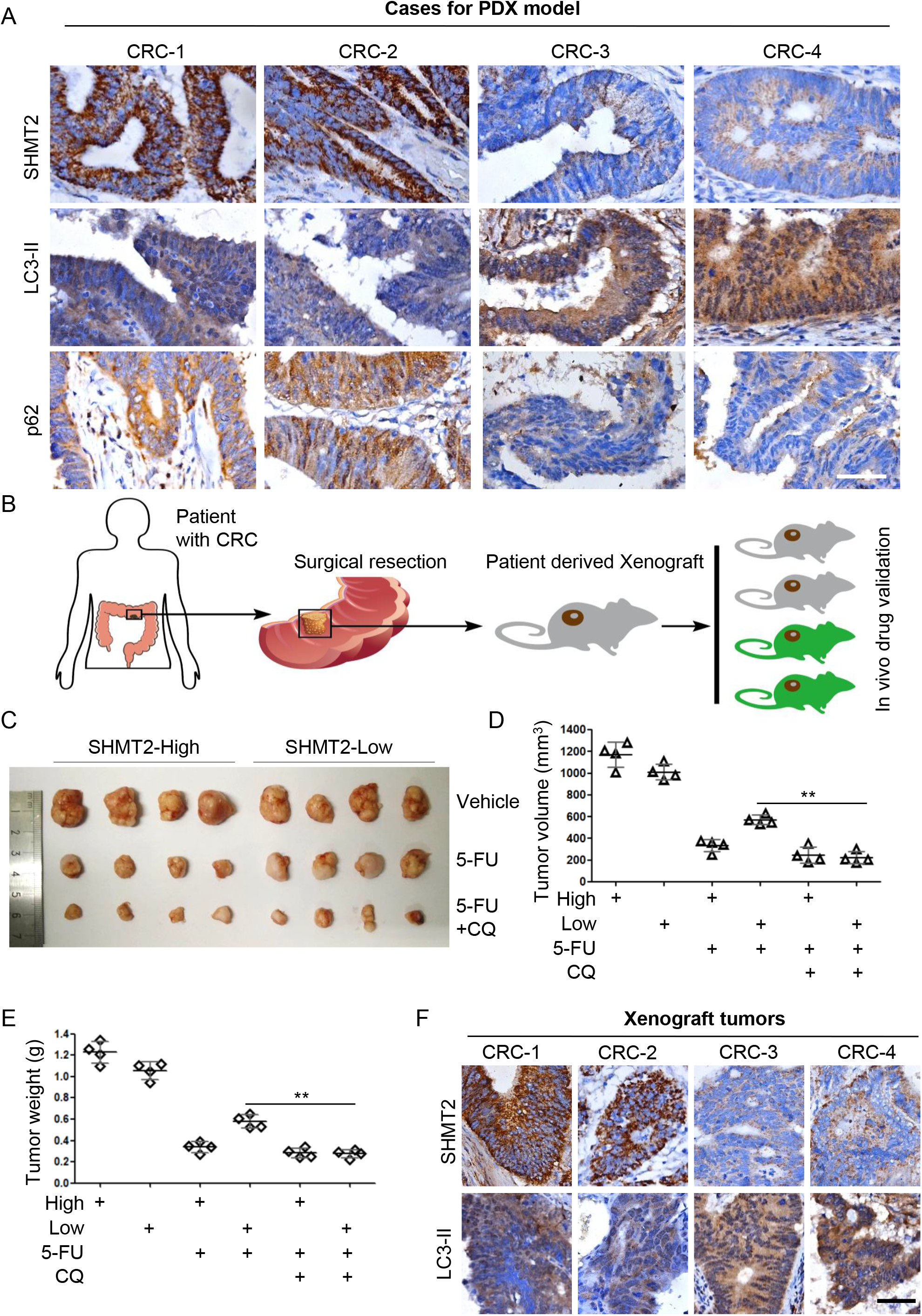
Chloroquine sensitized xenografts with low SHMT2 expression to 5-FU treatment in the PDX model. A Immunohistochemical images of SHMT2, LC3, and p62 staining in CRC tissues from four selected patients (two with low SHMT2 expression and two with high SHMT2 expression) using the indicated antibodies. B Schematic of the patient-derived xenograft (PDX) model. C-E The xenograft experiments with 5-FU or CQ treatment are described in the Methods section. (C) Xenograft tumours were harvested and photographed. The quantification of the (D) average volume and (E) weight of the xenograft tumours is shown. Four tumours from individual mice were included in each group, ** *P* < 0.01. F Sample immunohistochemical images of xenograft tumours. Scale bar, 50 μm.

## Discussion

We screened 66 differentially expressed genes associated with CRC progression in 224 colon cancer tissues and 165 adjacent normal tissues from three GEO data sets and found that SHMT2 is important in CRC metabolism. SHMT2, which is responsible for the conversion of serine to glycine, supports cancer cell proliferation in various cancers (5,15,33). SHMT2 is upregulated in CRC and plays an important role in colorectal carcinogenesis (6,8). Despite the genetic diversity of tumours, 37% of CRC cases showed low expression of SHMT2. However, SHMT2-low CRC showed 5-FU chemoresistance and poor prognosis. Further analysis revealed that SHMT2 induced autophagy and subsequently triggered 5-FU resistance. Using mass spectrometry, we identified cytoplasmic p53 as a SHMT2-binding protein and found that SHMT2 inhibited autophagy by stabilizing cytoplasmic p53. After 5-FU treatment, depletion of SHMT2 promoted autophagy and inhibited apoptosis. Inhibition of autophagy induced by low SHMT2 expression sensitized CRC cells to 5-FU treatment in vivo and in vitro. Last, we enhanced the lethality of 5-FU treatment to CRC cells through the autophagy inhibitor CQ in a patient-derived xenograft (PDX) model. These findings are important in terms of the response to 5-FU chemotherapy in SHMT2-low CRC cases (37%).

Autophagy has opposing and context-dependent roles in cancer, and therapeutic targeting of autophagy in cancer is sometimes viewed as controversial (21,22). In our study, we found that low expression of SHMT2 increased the resistance of CRC cells to 5-FU treatment through autophagy induction. The clinical data also verified this finding. In vivo and in vitro, depletion of SHMT2 induced 5-FU resistance, while autophagy inhibitors decreased this resistance. Thus, our study supports the hypothesis that autophagy inhibitors are beneficial to 5-FU-based chemotherapy in CRC.

The factors inducing autophagy are complicated (for example, starvation, rapamycin and toxins affecting the endoplasmic reticulum) (21,34). Among these factors, inhibition of p53 led to autophagy, and cytoplasmic p53 was able to repress the enhanced autophagy of p53^−/−^ cells. Some inducers of autophagy stimulate proteasome-mediated degradation of p53 through the E3 ubiquitin ligase HDM2. However, the factors regulating the binding of HDM2 to p53 need to be explored. In our study, we found that SHMT2 competitively bound cytoplasmic p53 to exclude HDM2 and thus inhibited autophagy. Treatment with 5-FU increased the binding of p53 to HDM2 to induce autophagy, while the binding of cytoplasmic p53 to SHMT2 decreased. In conclusion, we verified a new mechanism of the SHMT2-p53-HDM2 competitive binding system for the regulation of autophagy and confirmed the importance of the mechanism in CRC 5-FU-based chemotherapy.

SHMT2 is located not only in mitochondria but also in the cytoplasm, as shown by our data and those from other studies (9). SHMT2 is a tetrameric metabolic enzyme involved in one-carbon metabolism and can also participate in the BRISC-SHMT complex to deubiquitinate IFNAR1 and regulate interferon responses (9). In our study, SHMT2 regulated autophagy not by controlling one-carbon metabolism but by binding to cytoplasmic p53. These findings emphasize the multiformity of SHMT2 function and are helpful for the overall understanding of the function of SHMT2 in one-carbon metabolism and autophagy.

Given the complexity of tumourigenesis, mechanisms of chemotherapy still need to be explored. Regarding CRC chemotherapy, our study showed that low expression of SHMT2 is related to 5-FU resistance. These results imply that the expression of SHMT2 could be used as a marker for 5-FU resistance in clinical therapy. Exploration of the molecular mechanism showed that SHMT2 competitively binds cytoplasmic p53 to exclude the E3 ubiquitin ligase HDM2 and that depletion of SHMT2 decreases the stability of cytoplasmic p53 to induce autophagy, which maintains the survival of cancer cells treated with 5-FU. These findings reveal the binding of SHMT2 to p53 as a novel cancer therapeutic target and provide a potential opportunity to reduce 5-FU resistance using autophagy inhibitors in chemotherapy.

## Material and Methods

### Patients, cohorts and tissue microarrays

Fifty paired fresh frozen samples of primary colorectal carcinoma (CRC) and adjacent normal colon tissue were collected from the Department of Surgery at the Shanghai General Hospital, School of Medicine, Shanghai Jiaotong University. A total of 378 paraffin-embedded samples of stage II-III primary colorectal carcinoma were collected from 2003 to 2011 and made into tissue microarrays. All samples were obtained during surgery. This research was approved by the Ethics Committee of Shanghai General Hospital (2016KY069), and written informed consent was obtained from all patients before enrollment in the study.

### Immunohistochemistry

Polylysine-coated slides were deparaffinized in xylene and then rehydrated using a graded ethanol series. The slides were then put in 3% hydrogen peroxide to quench endogenous peroxidase activity, followed by heating in a microwave for 10 min in 10 mM citrate buffer (pH 6.0). Primary antibodies diluted in phosphate-buffered saline (PBS) containing 1% bovine serum albumin were added to the slides and incubated at 4°C overnight. Slides were then incubated with the secondary antibody (Genentech, Shanghai, China) for 30 min at room temperature, followed by counterstaining with Mayer’s hematoxylin. The immunohistochemistry slides were scanned by an Advanced CCD Imaging Spectrometer, which captured digital images of whole immunostained slides.

The intensity and extent of staining were evaluated independently by two pathologists who were blinded to patient outcomes. The staining intensity score was rated as 0 (no staining), 1 (mild staining), 2 (moderate staining), or 3 (intense staining). The extent score was assigned as 0 (0%), 1 (1%–25%), 2 (26%–50%), 3 (51%–75%), or 4 (76%–100%) according to the percentage of positively stained cells. The final scores were calculated by multiplying the intensity scores by the area scores. The patients with CRC were divided into two groups according to the staining score: 0–8, lower expression; 9–12, higher expression.

### Nude mouse xenograft models

CRC xenografts were established in 5-week-old female BALB/c nude mice purchased from the Institute of Zoology, Chinese Academy of Sciences of Shanghai. Briefly, SHMT2-KD and control cells (3 × 10^6^) were suspended in 100 μl of PBS and injected subcutaneously into both flanks of the 6-week-old nude mice. The tumour volume was calculated using the following formula: length × width2 × 0.5. When the tumour volumes were approximately 0.2 cm^3^, the mice were divided into four groups: Vehicle, 5-FU, 5-FU + CQ, and CQ. 5-FU was dissolved in PBS, and 20 mg/kg/day was injected intraperitoneally twice a week for three weeks. CQ (10 mg/kg) was administered as a daily oral gavage with a stainless steel ball-head feeding needle. The tumour volume and body weight of the mice were measured once a week, and the mice were sacrificed after five weeks. All of the animal procedures were conducted by the Hospital Animal Care guidelines of Shanghai Jiaotong University Affiliated Shanghai General Hospital. All efforts were made to minimize animal suffering.

### Establishment of patient-derived xenografts (PDX model)

Fresh tissues from 4 CRC patients (2 high-SHMT2 and two low-SHMT2) undergoing surgical treatment were obtained and transported immediately to the animal facility in PBS at 4°C. A primary tissue sample was anonymized and obtained by the Shanghai General Hospital (Shanghai, China) Institutional Review Board. Primary CRC tissue specimens were minced with scissors into small (2–3 mm^3^) fragments. Under aseptic conditions, tissue fragments were implanted subcutaneously into the flanks of female B-NSG mice (NOD-Prkdc^scid^ Il2rg^tm1^/Bcgen, Biocytogen) by using a 10-gauge trocar needle through a small incision on each animal’s dorsal flank. Animal health was monitored daily. Once established, solid tumour xenografts were serially passaged using the same technique.

### Cell lines, plasmids, and reagents

The human cell lines HCT116, SW480 and 293T were purchased from the American Type Culture Collection (ATCC, Manassas, VA, USA). All cell lines were maintained in DMEM supplemented with 10% FBS (Gibco, USA) at 37°C, 95% humidity and 5% CO2. The SHMT2, HDM2, p53 and p53 mutant (NLS-, NES-) sequences (29) were cloned into the pCDNA 3.0 vector or pLVX-IRES (lentiviral expression vector) vector by standard cloning methods. The pLKO.1-shRNAs targeting SHMT2 double-stranded oligonucleotides were as follows: 5’CCGG<u>ACAAGTACTCGGAGGGTTATC</u>CTCGAGGATAACCCTCCGAGTACTTGTTT TTTG (SHMT2-Sh-1), 5’CCGG<u>TAGGGCAAGAGCCAGGTATAG</u>CTCGAGTAGGGCAAGAGCCAGGTATAGT TTTTG (SHMT2-Sh-2), and 5’CCGG<u>GTCTGACGTCAAGCGGATATC</u>CTCGAGGATATCCGCTTGACGTCAGACTT TTTTG (SHMT2-Sh-3); the shRNA targeting p53 double-stranded oligonucleotides was as follows: 5’CCGG<u>CGGCGCACAGAGGAAGAGAAT</u>CTCGAGATTCTCTTCCTCTGTGCGCCGTTTTTG (p53-Sh) (target sequences are underlined). The Scramble shRNA targeting double-stranded oligonucleotides was 5’CCGG<u>CAACAAGATGAAGAGCACCAA</u>CTCGAGTTGGTGCTCTTCATCTTGTTGTTTTTG. The autophagy inhibitor 3-MA and CQ (chloroquine) were obtained from Sigma. Antibodies against LC3 (ABC929, Sigma; ab48394, Abcam), SHMT2 (NBP1-80755, Novus Biologicals USA), p62 (p0017, Sigma), p53 (DO-1, Santa Cruz; ab32389, Abcam) and actin (A1978, Sigma) were used.

### Western blotting and immunostaining

Cell lysate preparation and western blotting were performed as previously described (35).

### Quantification of GFP–LC3 puncta

The formation of autophagic vesicles was further monitored by assessing the transient expression of GFP-LC3 aggregates in HCT116 cell lines and was quantified by counting the percentage of cells showing accumulation of GFP–LC3 in dots or vacuoles (GFP–LC3^vac^) in a minimum of 100 cells per preparation in three independent experiments. Cells presenting a mostly diffuse distribution of GFP–LC3 in the cytoplasm and nucleus were considered nonautophagic, whereas cells presenting several intense punctate GFP–LC3 aggregates with no nuclear GFP–LC3 were classified as autophagic. Each GFP–LC3 staining sample was read by two independent investigators.

### Immunofluorescence

Immunofluorescence was performed as previously described (36).

### Gas Chromatography-Mass spectrometry

In total, six biological replicates of each cell group were subjected to GC/MS analysis. Briefly, 60 mg of each tissue section was transferred to a 1.5 ml tube. Metabolites were extracted by the addition of 360 μL of methanol (precooled to −20 °C) and 40 μL of internal standard (L-2- chlorophenylalanine), followed by vortexing for 2 min, sonication for 30 min, and, after the addition of 200 μL of chloroform and 400 μL of water, sonication again for 30 min. The derivatized samples were analyzed on an Agilent 7890A gas chromatography system coupled to an Agilent 5975C MSD system (Agilent, CA). An HP-5MS fused silica capillary column (30 m × 0.25 mm × 0.25 μm, Agilent J & W Scientific, Folsom, CA, USA) was utilized to separate the derivatives. Helium (> 99.999%) was used as the carrier gas at a constant flow rate of 6.0 mL/min through the column. The acquired MS data from GC−MS were analyzed by ChromaTOF software (v 4.34, LECO, St. Joseph, MI). Metabolites were qualitatively annotated by the Fiehn database, which was linked to the ChromaTOF software. The internal standard was used for data quality control (reproducibility). After internal standards and any known pseudo-positive peaks, such as peaks caused by noise, column bleeding and the BSTFA derivatization procedure, were removed from the data set, the peaks from the same metabolite were combined. The differential metabolites were selected on the basis of satisfying the combination of a statistically significant threshold of variable influence on projection (VIP) values obtained from the OPLS-DA model and *P* values from a two-tailed Student’s t-test on the normalized peak areas, where metabolites with VIP values greater than 1.0 and p values less than 0.05 were included.

### Statistical analysis

Student’s two-tailed t-test and GraphPad Prism were used for statistical analysis. Differences were considered significant at a p-value of < 0.05.

## Acknowledgments

This work was supported by National Natural Science Foundation of China (Grant nos. 81871931, 81602069, 81220108021, 81472241, 81670514, 81702337, 81602616), Medical Guidance Project of Shanghai Science and Technology Commission (Grant nos. 17411968200, 19QA1407100), Natural Science Foundation of Shanghai Municipality (Grant no. 16ZR1427700).

## Author contributions

JC, GF and ZP designed the study; GF, CX and XW performed bench experiments; YP, GS and JC conducted the animal experiments; HL and WG provided several PDX models; XL, HT and ZG performed the bioinformatics and statistical analyses; JC and ZP wrote the manuscript with comments from all authors.

